# Distinct profiles of tinnitus and hyperacusis in intensity deviant responses and auditory evoked potentials

**DOI:** 10.1101/2024.01.02.573726

**Authors:** Ekaterina A Yukhnovich, Kai Alter, William Sedley

**Affiliations:** Newcastle University

## Abstract

ERPs in response to intensity deviant stimuli are assessed in four age and hearing matched groups of various combinations of tinnitus and hyperacusis (both conditions, one of the conditions, neither condition). Distinct profiles for tinnitus and hyperacusis are shown, as well as additional more nuanced interactions. This not only moves our understanding of each condition, but also speaks directly to possible mechanistic subtypes of tinnitus (and of hyperacusis) which might be disentangled through the cheap and available technique that is single-channel EEG. The current findings may also explain some discrepant findings in past literature.

## Introduction

### Tinnitus & Hyperacusis

Tinnitus is a persistent sound heard by an individual without an environmental source [1]. It is a common condition, yet the search for a human biomarker of tinnitus is still on-going. There is a high correlation between presence of tinnitus and hyperacusis, which increases with severity of tinnitus [2, 3]. Hyperacusis is an auditory condition that causes normal environmental sounds to be uncomfortably loud [4]. It can strongly impact quality of life [5]. Hyperacusis is likely to occur with hearing loss and in the ageing population, though it also occurs in people with normal audiograms (possibly with hidden hearing loss such as asymmetric and notched audiometric results) [6–8]. In general, hyperacusis occurs due central gain changes, possibly from weakened inhibition from the auditory cortex to the inferior colliculus, and likely involves the limbic system [9, 10]. A previous fMRI study on adults with vs. without hyperacusis showed a global effect of enhanced sound-evoked activation across all frequencies [11], while another study showed reduced habituation (less N100 suppression) in patients with Fragile X syndrome who also had hyperacusis [12].

Four categories of hyperacusis have been described: loudness, annoyance/avoidance, fear, and pain [13]. Annoyance/avoidance and fear types are accompanied by strong emotions, while the last type is accompanied by pain around the face and the ear/s. Usually, the latter three types accompany the loudness hyperacusis, but have their own unique mechanisms [13]. Hyperacusis is highly prevalent in people with a variety of neurological conditions such as autism, ADHD, chronic pain, head traumas, depression, PTSD, etc [13], with some of these conditions being particularly related to fear and annoyance/avoidance hyperacusis (e.g. autism) while other are more related to pain hyperacusis (e.g. patients with chronic pain). Hyperacusis of an emotion-inducing subtype has also been related to chronic stress due to a connection between the amygdala and the autonomic nervous system. For example, rats did not adapt to prolonged exposure to noise and their levels of corticosterone were increased the longer they were exposed to this noise [14]. Another study showed a heightened connectivity between the orbitofrontal cortex/dorsal anterior cingulate cortex and the auditory cortex. Orbitofrontal cortex and dorsal anterior cingulate cortex have been previously implicated in pain perception and anticipation of noxious stimuli [13, 15]. People with hyperacusis also reported higher stress levels from their work environment compared to people without hyperacusis [16].

Much of the brain activity changes may be related to levels of hearing loss as well as distress caused by the tinnitus/hyperacusis [17]. Both tinnitus and hyperacusis may stem from abnormal central gain in the auditory cortex and related connections due to overcompensation by the brain for reduced environmental input, but in different ways [17, 18]. It has been suggested that tinnitus is the addition of central noise that compensates for reduced input by maintaining usual levels of activity, possibly through prediction errors, by increased spontaneous firing rates and/or heightened neural synchrony. On the other hand, hyperacusis stems from multiplicative central gain [19, 20]. Another theory states that a larger extent of peripheral pathology/deafferentation may lead to a failure to compensate sufficiently by increasing central gain, which is related to tinnitus, while hyperacusis may relate to the over-amplified compensating central gain increase [7]. Both processes could occur in one person, from which an interaction between the two conditions could be seen that could in turn alter the shape of studied activity.

However, despite so much overlap between tinnitus and hyperacusis prevalences and aetiologies, much of earlier research did not account for or focus on hyperacusis as a potential confounding factor. Hyperacusis presence may affect patterns and regions implicated in tinnitus research, though the existence of a connection between the two conditions has been long-known [19, 21, 22]. As an example of a problem with such conflations in previous research, an fMRI study showed that activation in inferior colliculus and medial geniculate body, previously correlated with tinnitus, was more directly related to reactivity to external sounds, which was not accounted for in other studies [18]. This oversight has come to light in the recent years, along with the realisation that many hyperacusis-focused studies mainly test subjects with minimal hearing loss [11]. Some more recent studies focus on comparisons of activity in tinnitus with and without hyperacusis [11], for example, an fMRI study demonstrated that responses to auditory stimuli in subcortical auditory structures and cortices (such as inferior colliculus and medial geniculate body) are increased when participants have hyperacusis (as well as overt moderate hearing loss and tinnitus). This finding was not due to reduced cortical frequency specificity that could be seen in the presence of hearing loss. Interestingly, they also established that the group with both tinnitus and hyperacusis had significantly smaller activation in response to tinnitus-like frequency stimuli compared to the group without hyperacusis but with tinnitus. There are two potential explanations for this phenomenon. The hyperacusis group may have more central noise at the tinnitus frequency, meaning that less gain is needed. Alternatively, this could occur due to ceiling effect of the fMRI-related background noise: tinnitus frequencies are already hyper-stimulated, so they have less room to increase their activity even further when played a matched tone [23].

#### Event-related potentials

Auditory event-related potentials, such as P50, N100, and Mismatch Negativity (MMN) have been used to investigate mechanisms and as potential biomarkers for a number of conditions, including tinnitus, schizophrenia, autism, mild cognitive impairment, Alzheimer’s disease and psychosis [24–29].

#### Standard responses

Generally, P50, N100 and P200 components are related to an inhibitory function called sensory gating, such as reducing responses to repetitive stimuli (filtering out irrelevant stimuli), or detecting novel stimuli [30]. It is a multistage mechanism, with each of the three ERPs having their own function [31]. P50 and N100 usually represent a pre-attentional and an attention-triggering filter mechanism, occurring around 50 ms and 100 ms after stimulus onset, respectively [32]. P200 may be related to allocation of attention, and can be an independent component from the N100 despite earlier research possibly suggesting otherwise [33]. These ERPs decrease in amplitude as a result of repeated stimuli. A reduction in P50 and N100 amplitudes reflect sensory gating-out of a repeating stimulus. One study showed that while change in frequency did not cause significant increase in P50 or N100, whereas intensity change did [30]. P200 has also been found to be intensity dependent [34].

Attenuated reduction in amplitudes to a second stimulus compared to an identical first stimulus have been found to be indicative of altered information processing in conditions such as schizophrenia and even in children with cochlear implants [31, 35]. Further, in a study using 1 kHz tone bursts, a bilateral tinnitus group were more sensitive to an increase in intensity from 70 dB to 90 dB SPL in terms of increasing N100 and P200 amplitudes, compared to unilateral tinnitus and control groups. However, inhibitory mechanisms are also diminished in hearing loss, so any conclusions must be done carefully [36].

N100 has also been implicated in the stimulus-specific adaption theory [37]. One study used trains of tones with a number of potential deviants. A standard repeating pattern included two tones (A B), one was a lower frequency tone (800 Hz) (A) which was then followed by a tone of a higher (1600 Hz) (B) frequency. Two of the possible deviants from this pattern had the first frequency repeat as the second tone (A A or B B), instead of the usual higher frequency tone. N100 suppression was larger to the deviants AA and AB compared to the standard AB, showing that N100 adapts to the physical attributes of the stimuli rather than more complex irregularities. On the other hand, MMN responses were larger to the deviants AA and AB compared to standard AB, thus showing the more complex processing underlying MMN.

In a study with 15 mild-tinnitus participants with normal hearing in normal and extended frequency audiometry testing, paired stimuli were presented to test sensory gating function [38]. Compared to the age and hearing-matched control group, the tinnitus group showed no significant differences in the gating processes, however, within the tinnitus group, there were ‘high suppressors’ and ‘low suppressors’. The two suppressor groups were defined based on the Pa amplitude, a small positive peak just before the P50 response. The high suppressors showed similar activity to the control group, but low suppressors increased early P50 amplitude to the second tone, as well as overall reduced N100 to both tones. This group also had an overall higher P200. Notably, hyperacusis was not measured in this study, therefore it is not possible to assess whether any differences between the tinnitus groups were due to hyperacusis presence or a different unmeasured factor. A review showed that N100 finding have been inconsistent so far both in the matching of hearing loss of tinnitus and control group, in the findings themselves both at frequencies below or within the tinnitus frequency of a participant [39]. Studies investigating solely hyperacusis are scarce [40], however, MEG studies on children with autism showed that M100 latencies were longer in children with auditory sensitivity compared to children without auditory sensitivity, and M50 dipoles were larger [41, 42]. In a group of children with Williams syndrome and hyperacusis, amplitudes in the P50-N100-P200 complex and MMN were increased compared to controls [43]. A study that used pairs of identical stimuli, found that P50 amplitude could predict acceptable noise level, which is associated with sensory gating functioning [44].

### Mismatch Negativity

MMN is a neural correlate of change detection [37]. Within the predicting coding framework, MMN occurs when a comparison between an expected stimulus and the sensory input do not match, thus producing a prediction error. The larger the difference, the larger the prediction error. Another theory of MMN is that it reflects stimulus-specific adaptation based on sensory memory, where repetition suppression occurs after a stimulus is repeatedly presented. In an fMRI study, it was found that MMN contains a number of subcomponents that stem from different cortical regions, which differentiate the responsibility for simpler feature detection (similarly to N100) and more complex processes such as prediction violation and uncertainty [37].

### Previous MMN research in tinnitus

An MMN-based biomarker in humans, called Intensity Mismatch Asymmetry (IMA), was identified as a potential tool for identifying a tinnitus biomarker when measured in response to intensity deviants at tinnitus frequency. IMA was based in predictive coding, aimed to relate to mechanisms forming part of a ‘final common pathway’ for tinnitus, irrespective of specific contributory mechanisms [24, 45]. Such a biomarker would help to better understand tinnitus mechanisms and allow treatment studies to determine the effectiveness of their treatment across tinnitus groups [20, 46]. IMA showed that participants with tinnitus had larger MMN responses to upward deviants (UD), but smaller MMN responses to downward deviants (DD), compared to the control group [24]. This first study focused on frequencies close to the tinnitus frequency only, and therefore a second study was conducted in which the tinnitus frequency responses were compared to responses to a 1 kHz control frequency [25]. However, the second study did not find the same pattern of activity. Potential reasons could have been higher levels of hearing loss overall, the overall paradigm context (the addition of much lower frequency), but also presence of hyperacusis. It is not known whether participants had hyperacusis in the original study, however in the second study, the majority of the tinnitus group had hyperacusis based on the updated cut-off score of 16 on the Khalfa Hyperacusis Questionnaire [47, 48].

Both tinnitus and/or hyperacusis have different effects on brain function, including MMN (Mismatch Negativity) amplitude [6, 19, 24, 49]. For example, when comparing MMN responses in a multi-deviant paradigm, tinnitus participants with normal extended frequency hearing had weaker MMNs for frequency, intensity, duration, location and silent gap deviants, compared to a age and gender matched control group [50]. The tones involved in this study were 500-1500 Hz, therefore far away from the usual tinnitus frequencies. Hyperacusis was not measured. In another example of a study that utilised a non-standard multi-feature MMN paradigm that included noise, pitch, location, intensity, laterality and rhythm deviants, with standards being chords that could often be heard in Western music, researchers found that central auditory mechanisms are altered in subjects depending on whether they had low/medium/high noise sensitivity (but no tinnitus) [49]. As tinnitus and hyperacusis can affect similar processes in different (or similar) ways, disentangling the two conditions is an important step in furthering our understanding of both conditions. Therefore, it would be useful to understand how each condition affects the auditory event-related components such as P50, N100, P200 and MMN, especially when these components are often used as a measure of tinnitus.

There are numerous motivating factor for the present study, which is a rigorous and systematic exploration of hyperacusis, tinnitus, and their combination, on early and late evoked responses and intensity deviant-related MMN. To my knowledge, this is the first study to assess ERPs between all four combinations of tinnitus and hyperacusis (both conditions, one of the conditions, neither condition), as well as the first study to investigate differences between responses to deviant stimuli overall, and specifically intensity deviant stimuli, between these groups. The four groups in this study are controlled for hearing profile, eliminating hearing loss as a potential confound. This should allow us to reflect on whether any of the previous tinnitus research tells us more about the involvement of hyperacusis than tinnitus specifically.

## Materials & Methods

### Participants

Four groups, each of 21 participants, were studied: 1) tinnitus without hyperacusis (T+H-), 2) tinnitus and hyperacusis (T+H+), 3) no tinnitus and no hyperacusis (C) and 4) hyperacusis without tinnitus (T-H+). Participants were recruited from affiliated volunteer lists at Newcastle University and via Google Ads. General inclusion criteria included being over 18 years old, and able to make an informed choice about volunteering. General exclusion criteria included using ongoing sedating or nerve-acting medications, and mental health conditions severe enough to interfere with everyday life activities. All potential participants needed to complete the Hyperacusis Questionnaire (HQ) [47] to understand which groups of participants they belonged to and whether they were eligible for the study. Those with an HQ score under 16 were in the H+ groups [48]. Those who scored above this, and also scored above 56 on the Inventory of Hyperacusis Symptoms (IHS) [51] were included in the H+ groups. To be included in one of the T+ groups, inclusion criteria also involved having chronic tinnitus for over 6 months, without an objective physical source, and which was not due to Meniere’s disease (this was an exclusion criterion for T-groups). T-participants were individually matched to T+ participants, based on an approximate match of their overall audiometric profiles, with particular attention to the vicinities of 1 kHz and the tinnitus frequency. It was also ensured that there were no significant group differences between equivalent tinnitus and control groups in age or sex.

Approval was given by the Newcastle University Research Ethics Committee, and all participants gave written informed consent according to the Declaration of Helsinki (reference number 5619/2020).

### Common methods: tinnitus psychophysics and EEG

The psychophysical assessment in which the tinnitus frequency and the intensity of sounds played during the EEG recording were determined, and the experimental design, followed the procedure in [25]. EEG data processing followed the procedure in [25], with the exception that rather than Denoising Source Separation, Independent Component Analysis was used to remove ocular artefacts.

### Statistical analysis

Statistical analysis was performed using MATLAB. To compare the evoked responses in participants with and without tinnitus and hyperacusis, three-way ANOVAs were used, with subject group (T+H-, T+H+, C, T-H+), frequency, and intensity used as factors of interest, and including interaction terms. Post-hoc analysis included Tukey Honest Significance Tests to determine any significant differences between the ERP amplitudes of the four groups, as this test is powerful when testing multiple numbers of means, and uses a similar parameter to the ANOVA tests [52]. Then, visual inspection of error bars created for the results section was carried out, based on which potential paired t-tests were run to see any patterns within-group, e.g. to see significant differences between downward deviant and upward deviant conditions.

## Results

### Demographic information

Table 1 shows means and standard errors (SE) of the demographic information of the 4 groups, their HQ scores, and THI scores in the two groups with tinnitus. One-way ANOVA showed that age was not significantly different between the four groups (p=0.892). HQ, on the other hand, was significantly different between the 4 groups (p<0.001). T+H+ and T-H+ had no significant differences in their HQ scores (p=0.089), whereas both H+ groups had significantly higher HQ scores than both C and T+H-groups (all p<0.001). This was expected as this was one of the bases for allocating groups. The cut-off score for HQ was 16 therefore all 4 groups were comfortably within their hyperacusis presence requirements. Following previous literature, the sample with tinnitus and hyperacusis had higher THI scores than those with tinnitus only. Chi-square test was carried out to establish that there was no significant difference in gender across the four groups (Chi (3) =2.61, p=0.456). As the pure tone audiometry results were not normally distributed at the majority of frequencies/ears Kruskal-Wallis tests were performed and showed that across all 4 groups, there were no significant differences in the hearing ability at 0.25, 0.5, 1, 2, 4, 6 or 8 kHz in right or left ears, with the smallest p-value being p=0.163.

**Table 1.** Descriptive statistics of the four study groups. Means and standard errors are given for every group for their age, HQ scores and THI scores. The gender split is also indicated for each group.

|  | T+H- | T+H+ | C | T-H+ |
| --- | --- | --- | --- | --- |
| Age | 53.71 (3.29) | 55.24 (3.17) | 54.33 (3.40) | 56.95 (2.47) |
| Gender | 13 f, 8 m | 13 f, 8 m | 11 f, 10 m | 16 f, 5 m |
| HQ scores | 10.24 (0.74) | 25.76 (1.35) | 5.29 (0.91) | 30.71 (1.61) |
| THI scores | 20.29 (3.13) | 45.52 (5.43) | N/A | N/A |

### Time course of the stimulus response

Grand average ERP data for channel FCz (with P9/P10 reference) across standard and deviant responses for all stimulus conditions and in each group was used to determine timeframes for quantifying P50, N100, P200 and MMN responses, based on visual inspection (Fig 1). To calculate the MMN difference waveform, standard responses were subtracted from their equivalent deviant conditions (Fig 1).

**Fig 1.**
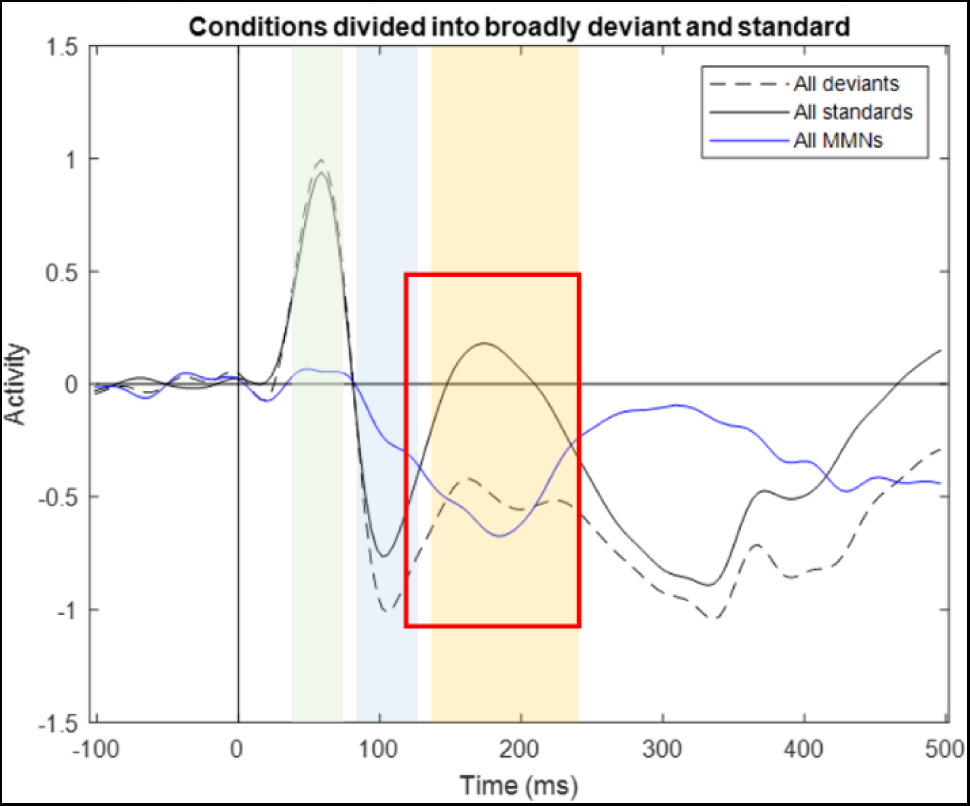
Standard, deviant and MMN waveforms broadly combined across groups and conditions. The chosen P50 timeframe was 45-75 ms (green), N100 was 85-120 ms (blue), P200 was 140-240 ms (orange). The MMN timeframe was 115-240 ms (red square). As the MMN and P200 overlap, P200 was only used in standards (and MMN in the difference waveform).

### Standard and pure deviant waveforms

The average waveforms to each condition are shown in Fig 2.

**Fig 2.**
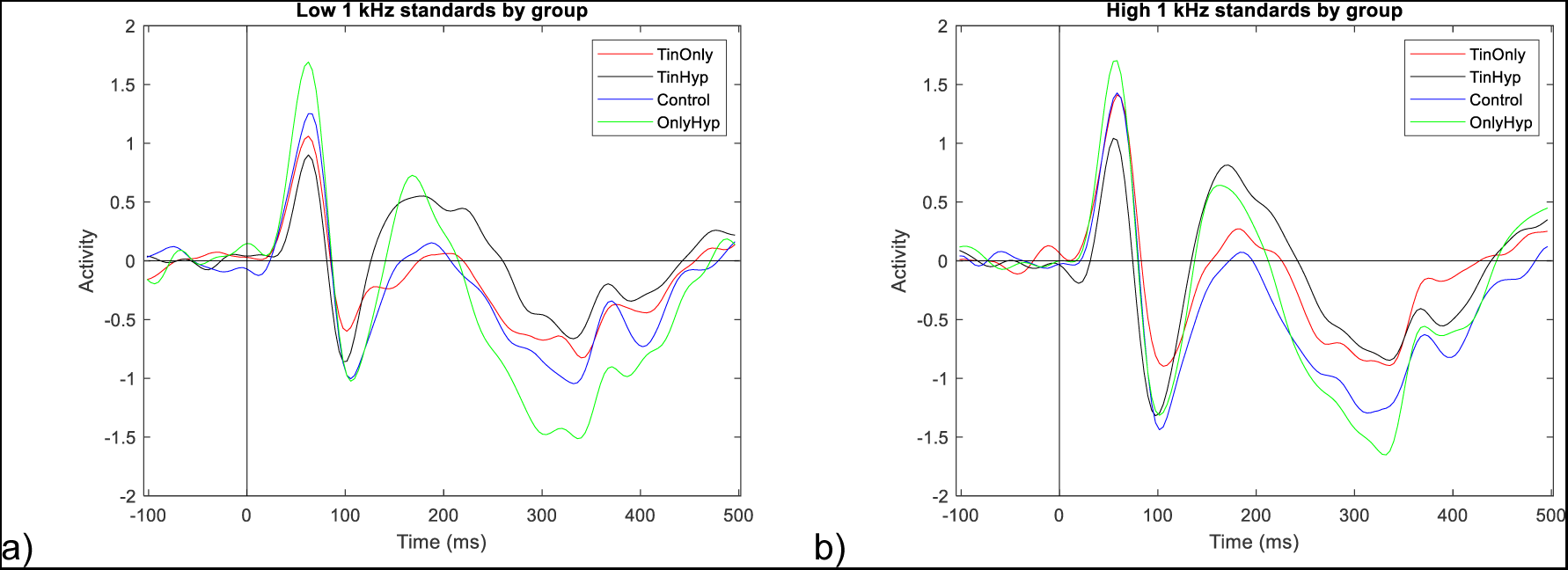
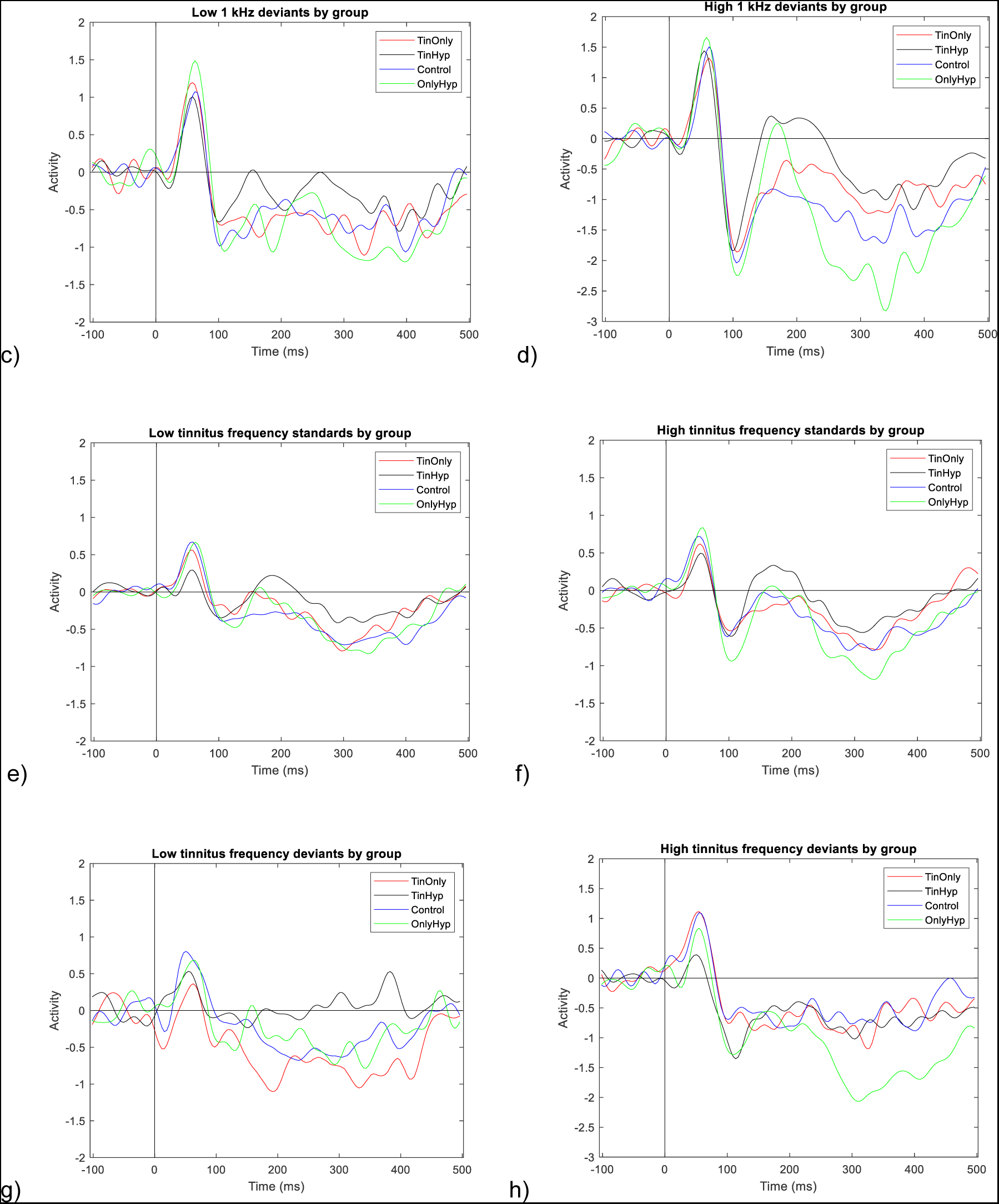
Average waveforms to standard and deviant conditions. Graphs a-d show responses at 1 kHz frequency, and e-h show responses to the tinnitus frequency. On the left are the quieter stimulus conditions and on the right are the louder stimulus conditions. The groups are colour-coded as follows: T+H-(red), T+H+ (grey). C (blue), T-H+ (green).

### P50 responses

#### Standards

Fig 3 shows P50 responses to standard stimuli in the four conditions (two intensities at two frequencies). A three-way ANOVA (group, frequency, intensity) showed significant main effects of group (p<0.001) and frequency (p<0.001), as well as an interaction between these two factors (p=0.011). Post-hoc test on group effects was carried out using Tukey test.

**Fig 3.**
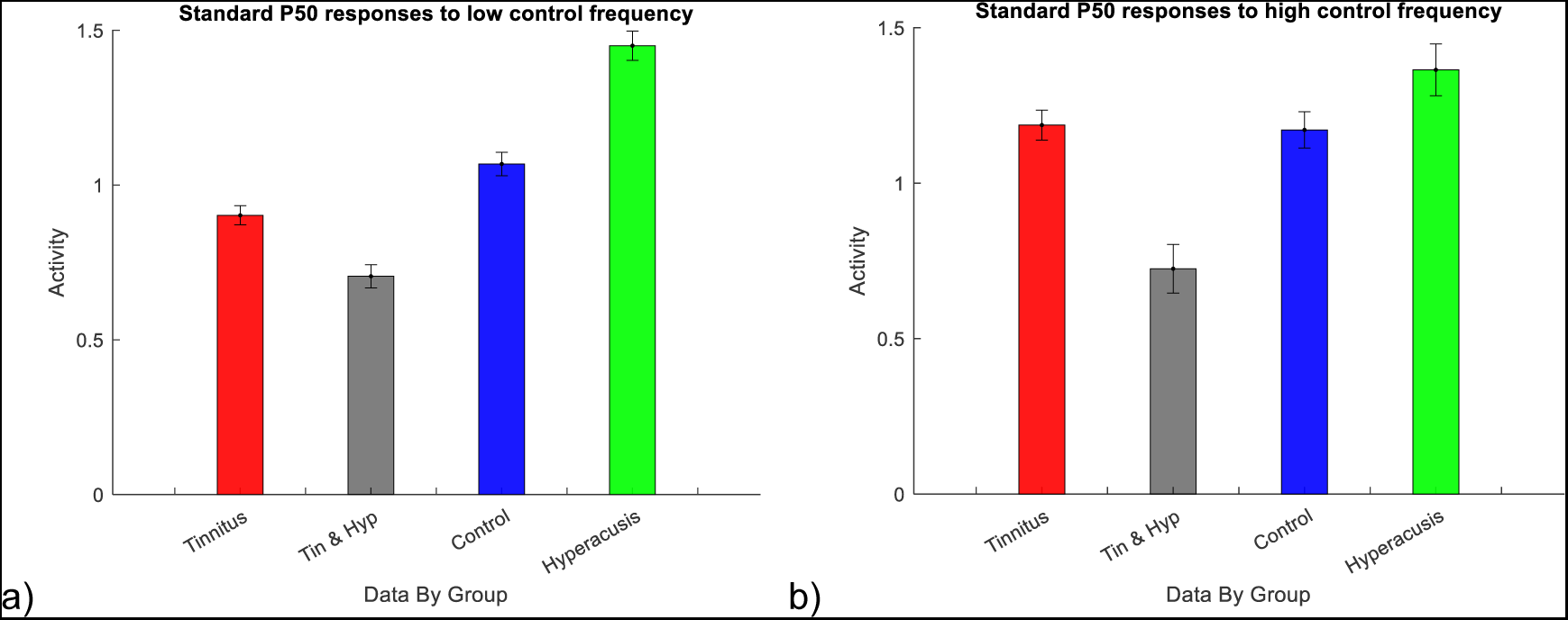
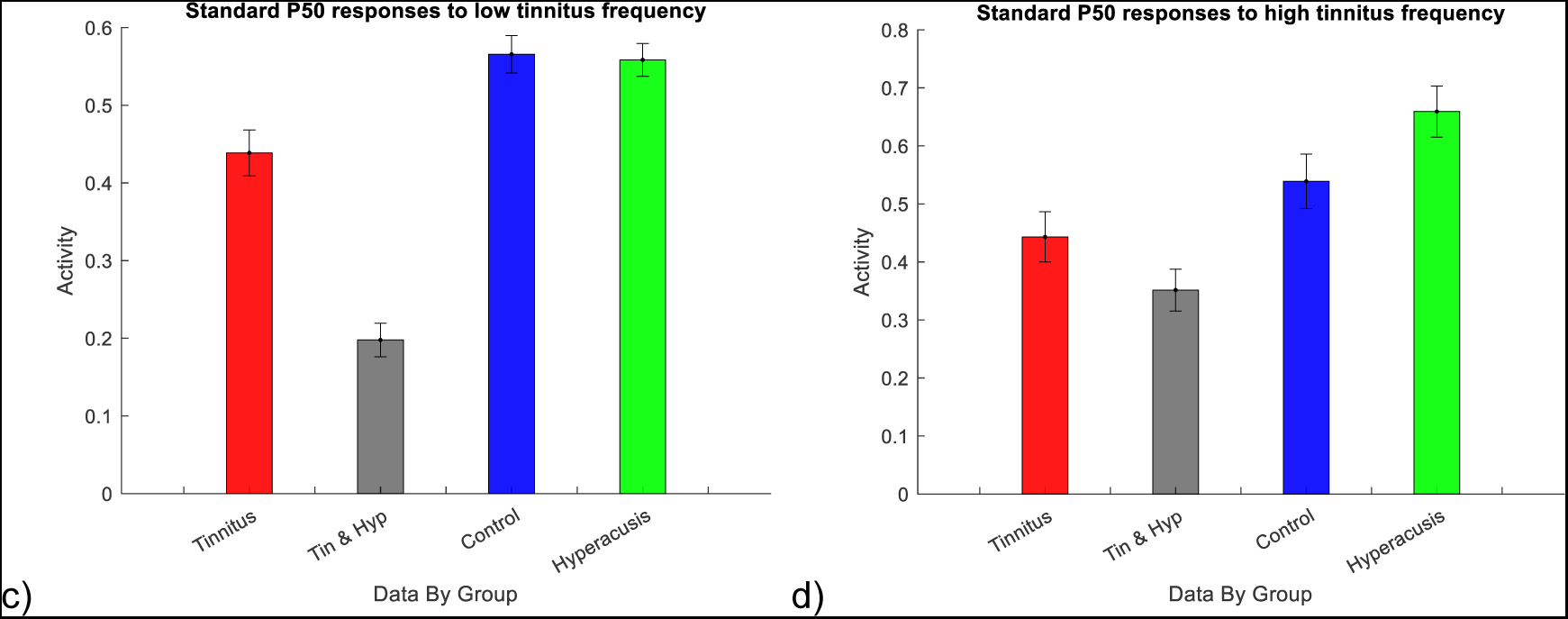
Standard responses in the P50 timeframe. Top two charts represent averaged responses to stimuli at 1 kHz frequency and bottom charts represent averaged responses to stimuli at tinnitus frequency. The order of the groups in all charts is: T+H-(red), T+H+ (grey). C (blue), T-H+ (green).

At low intensity stimuli at 1 kHz frequency, T-H+ group had a significantly higher amplitude than all other groups (all p<0.001); C group also showed significantly stronger response than T+H+ group (p=0.002); there were no significant differences between T+H- and T+H+ (p=0.146) or T+H- and C (p=0.266) groups. At high intensity at 1 kHz frequency, T+H+ group had significantly smaller amplitude compared to all other groups (all p<0.001); no other significant differences were seen.

At low intensity at tinnitus frequency, T+H+ group had significantly lower P50 amplitude compared to T+H- (p=0.031), control (p=0.039) and T-H+ (p=0.002) groups. There were no significant differences between T+H- and C groups (p=0.999), or C and T-H+ groups (p=0.613). At high intensity at tinnitus frequency, T+H+ group had significantly lower amplitude than T-H+ group (p=0.020), no other significant differences were seen.

#### Deviants

Fig 4 shows P50 responses to raw deviant (not difference waveform) stimuli in the four conditions. A three-way ANOVA (group, frequency, intensity) was carried out. All factors showed a significant main effect (all at p <.001). There was a significant interaction between intensity and group (p=0.004) affecting p50 deviant amplitudes. There was also a significant interaction effect of group and frequency (p<0.001), as well as an intensity, group, frequency interaction effect (p<0.001). Post-hoc Tukey tests were carried.

**Fig 4.**
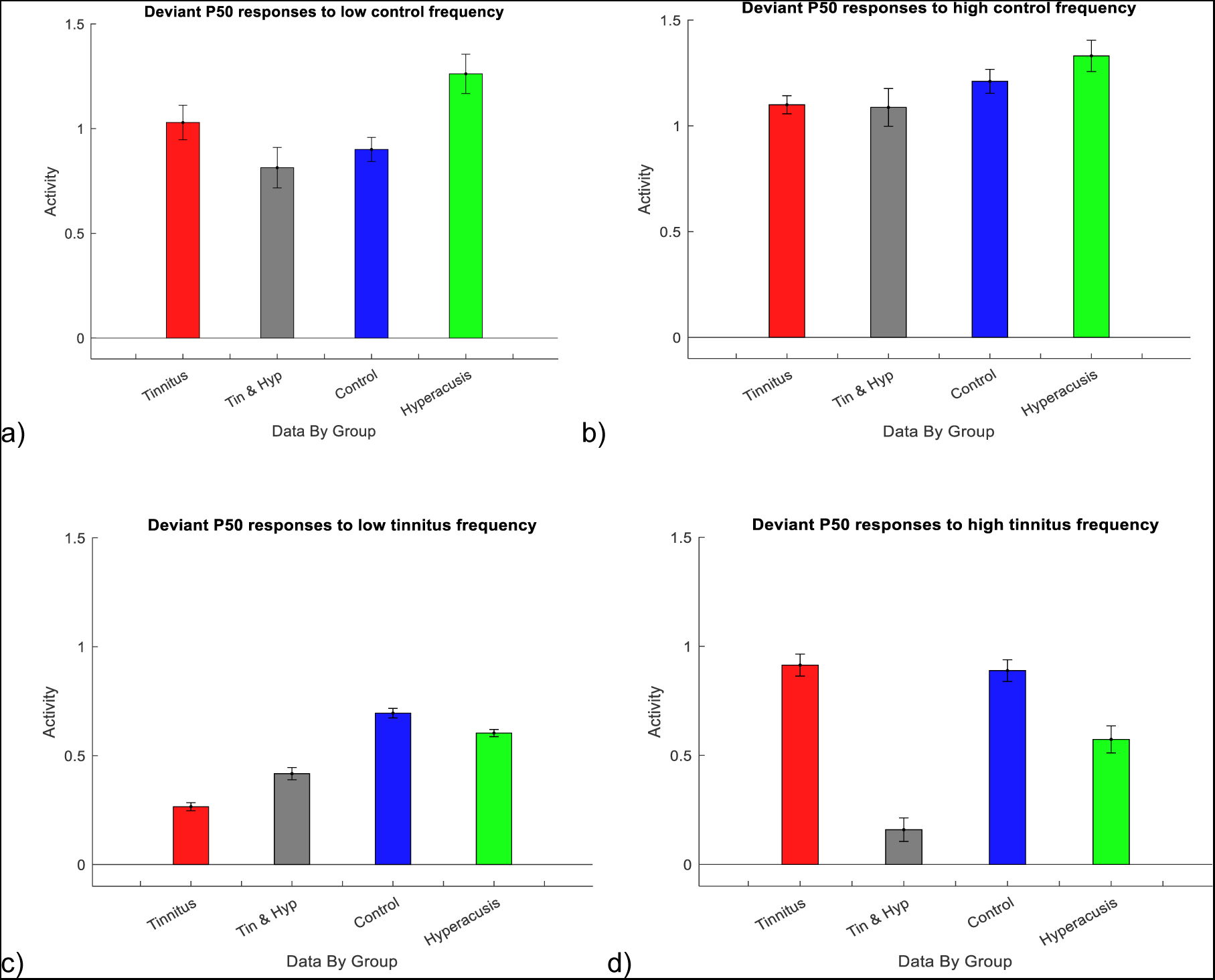
Deviant responses in the P50 timeframe. Top two charts represent averaged responses to stimuli at 1 kHz frequency and bottom charts represent averaged responses to stimuli at tinnitus frequency. The order of the groups in all charts is: T+H- (red), T+H+ (grey). C (blue), T-H+ (green).

At low intensity 1 kHz deviant, T-H+ group had significantly higher amplitude compared to C group (p=0.003) and T+H+ group (p<0.001); there were no other significant differences. At high intensity 1 kHz deviant, there were no significant differences.

At low intensity tinnitus deviant, T+H- group had significantly lower amplitude compared to C and T-H+ groups (both p<0.001) as well as T+H+ group (p=0.024); T+H+ group also had significant lower amplitude compared to C group (p<0.001) and T-H+ group (p=0.004); there was no significant difference between C and T-H+ group (p=0.275). At high intensity tinnitus deviant, T+H+ group had significantly lower amplitude compared to T+H- and C groups (both p<0.001) as well as T-H+ group (p=0.012).

Based on observation of Fig 4 and ANOVA results, post-hoc within-subject tests were carried out, to investigate whether intensity at tinnitus frequency significantly affects the amplitude of p50 responses in the two T+ groups. When responses to low intensity were compared to high intensity in the tinnitus frequency, both T+ groups showed significant differences but in opposite directions. The T+H- group had significantly higher amplitudes in response to a high intensity deviant (t(7)=-8.45,p<0.001), while T+H+ group had lower amplitudes in response to a high intensity deviant (t(7)=5.59,p<0.001).

### N100 responses

#### Standards

Fig 5 shows the mean amplitudes of N100 responses to standard tones in the four conditions. A three-way ANOVA (group, stimulus frequency, stimulus intensity) showed main effects of the three factors (all p<0.001). There were interaction effects of group and frequency (p=0.004), and a non-significant trend of group, frequency and intensity interaction (p=0.054). Tukey post-hoc analysis was carried out at the two frequencies.

**Fig 5.**
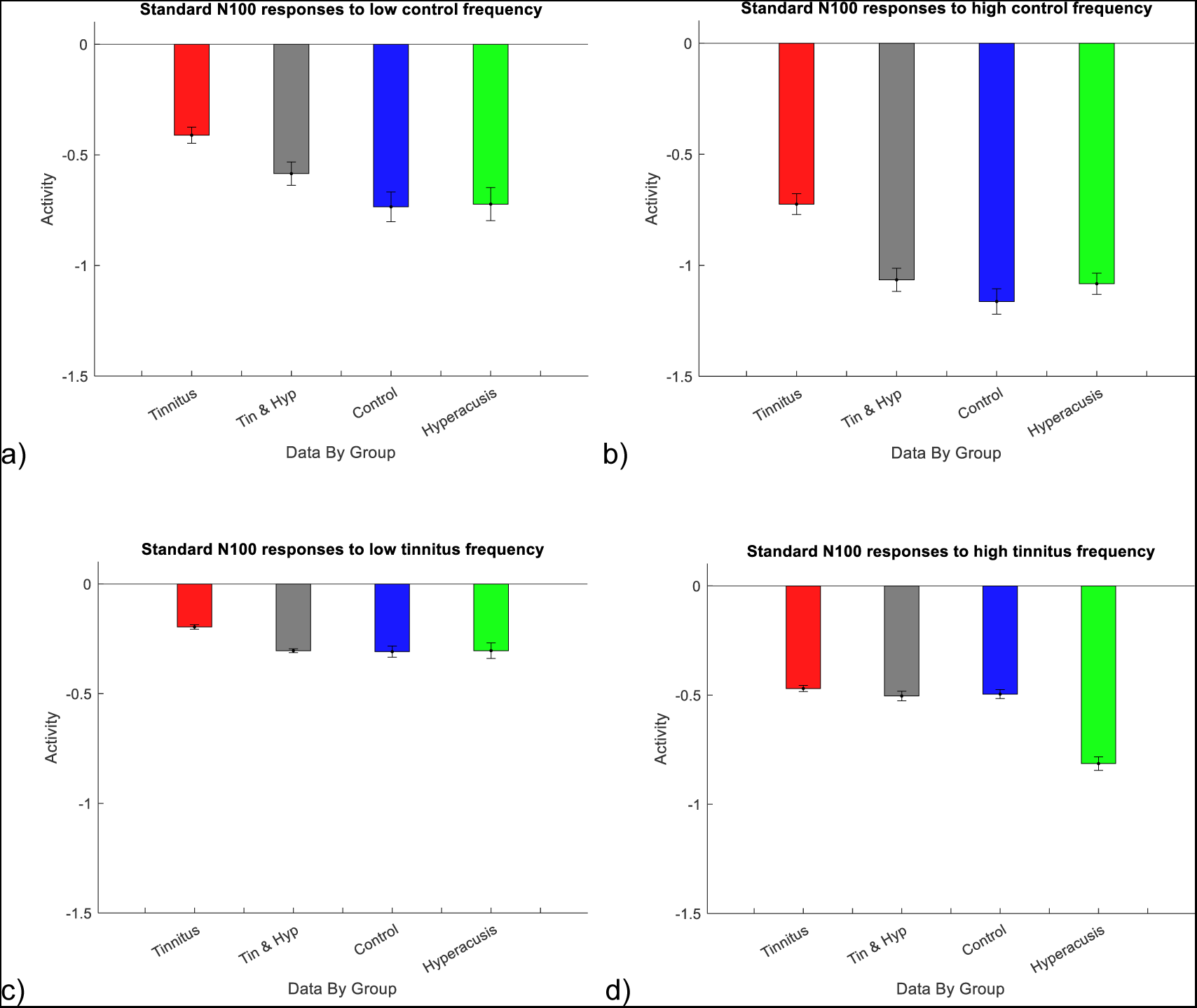
Standard responses in the N100 timeframe. Top two charts represent averaged responses to stimuli at 1 kHz frequency and bottom charts represent averaged responses to stimuli at tinnitus frequency. The order of the groups in all charts is: T+H- (red), T+H+ (grey). C (blue), T-H+ (green).

At low intensity 1 kHz frequency, there were no significant differences between groups. At high intensity 1 kHz frequency, significantly weaker amplitude was seen in T+H- groups compared to T+H+ (p=0.013) and C (p<0.001) and T-H+ (p=0.008) groups; no other significant differences were present.

At low intensity tinnitus frequency, there were no significant differences between the groups. At high intensity tinnitus frequency, T-H+ group had significantly stronger amplitude compared to all other groups (all p<0.001); no other significant differences were present.

#### Deviants

Fig 6 shows the mean amplitudes of N100 responses to deviant tones in the four conditions. A three-way ANOVA (group, frequency, intensity) showed main effects of all factors (p<0.001 for all), as well as interaction effects between group and frequency (p=0.001), group and intensity (p=0.001), and frequency and intensity (p<0.001). There was also a significant interaction effect between group, frequency and intensity (p=0.022). Tukey post-hoc analysis was carried out at the two frequencies.

**Fig 6.**
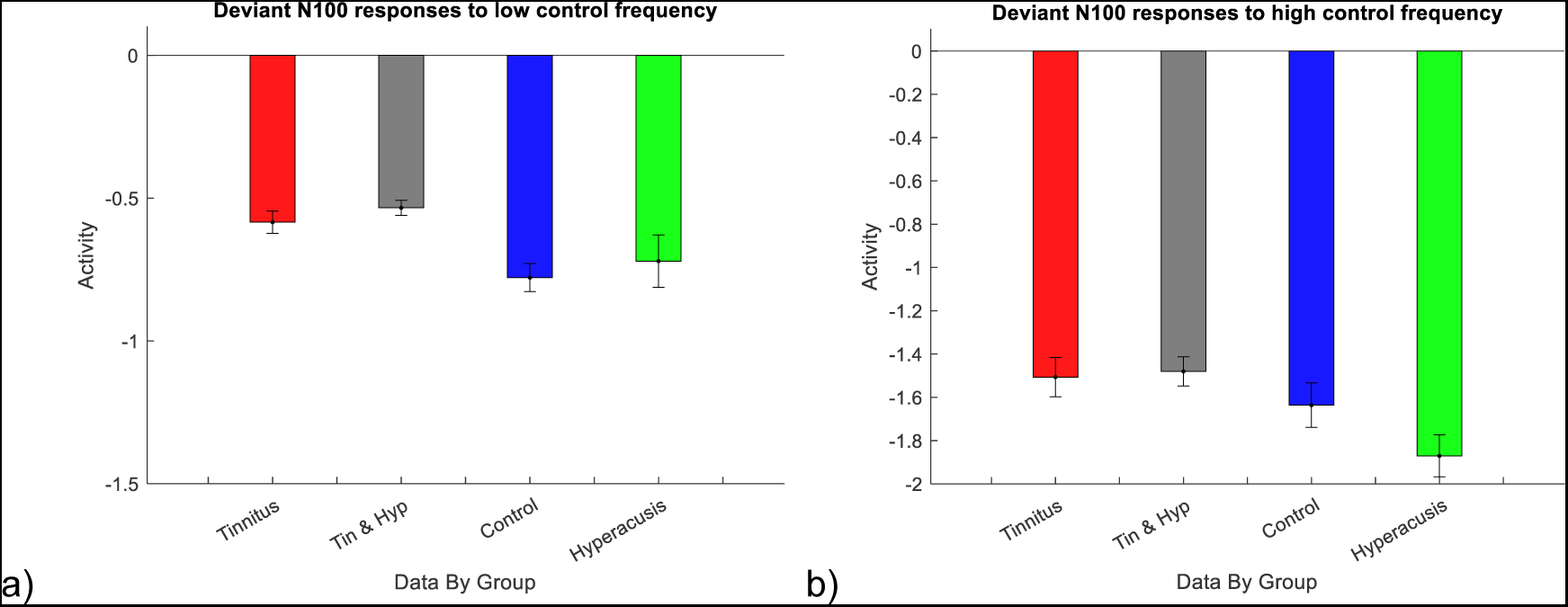
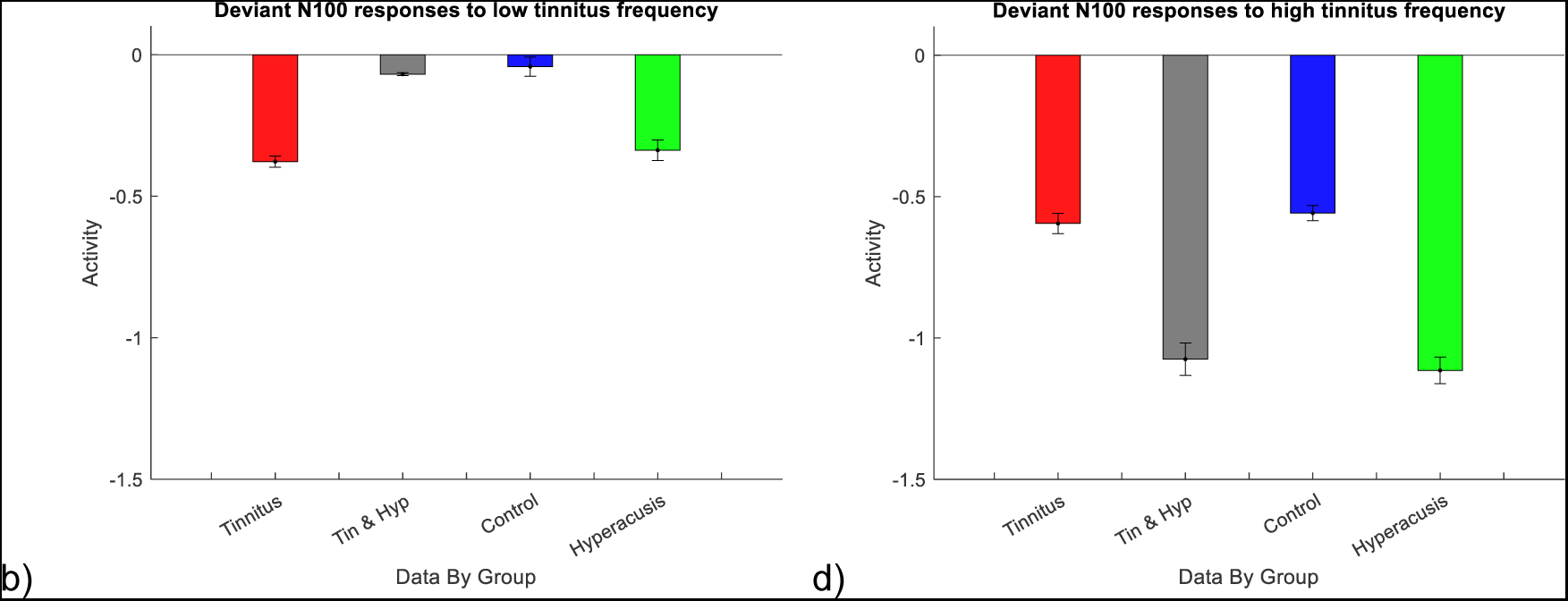
Deviant responses in the N100 timeframe. Top two charts represent averaged responses to stimuli at 1 kHz frequency and bottom charts represent averaged responses to stimuli at tinnitus frequency. The order of the groups in all charts is: T+H- (red), T+H+ (grey). C (blue), T-H+ (green).

At both low and high intensity 1 kHz deviants, no significant differences were seen between the groups.

At low intensity tinnitus deviant, significantly stronger responses were seen in T+H- group compared to T+H+ and control groups (both p<0.001); also, both T+H+ and C groups had significantly weaker responses compared to T-H+ group (both p<0.001). At high intensity tinnitus deviant, however, T+H- group had weaker responses than T+H+ and hyperacusis only groups (both p<0.001); T+H+ group had significantly higher responses than C group (p<0.001); no significant differences were seen between T+H- and C groups (p=0.976) or between T+H+ and T-H+ groups (p=0.970).

Post-hoc, paired t-tests were carried, that showed significant differences in responses to the two intensities within each group (all p<0.001).

### P200 responses

#### Standards

Fig 7 shows the mean amplitudes of P200 responses to standard tones in the four conditions. A three-way ANOVA (group, frequency, intensity) showed main effects of group and frequency (p<0.001) but not intensity (p=0.173). There were interaction effects between group and frequency, and group, intensity and frequency (both p<0.001). Post-hoc Tukey tests were carried out.

**Fig 7.**
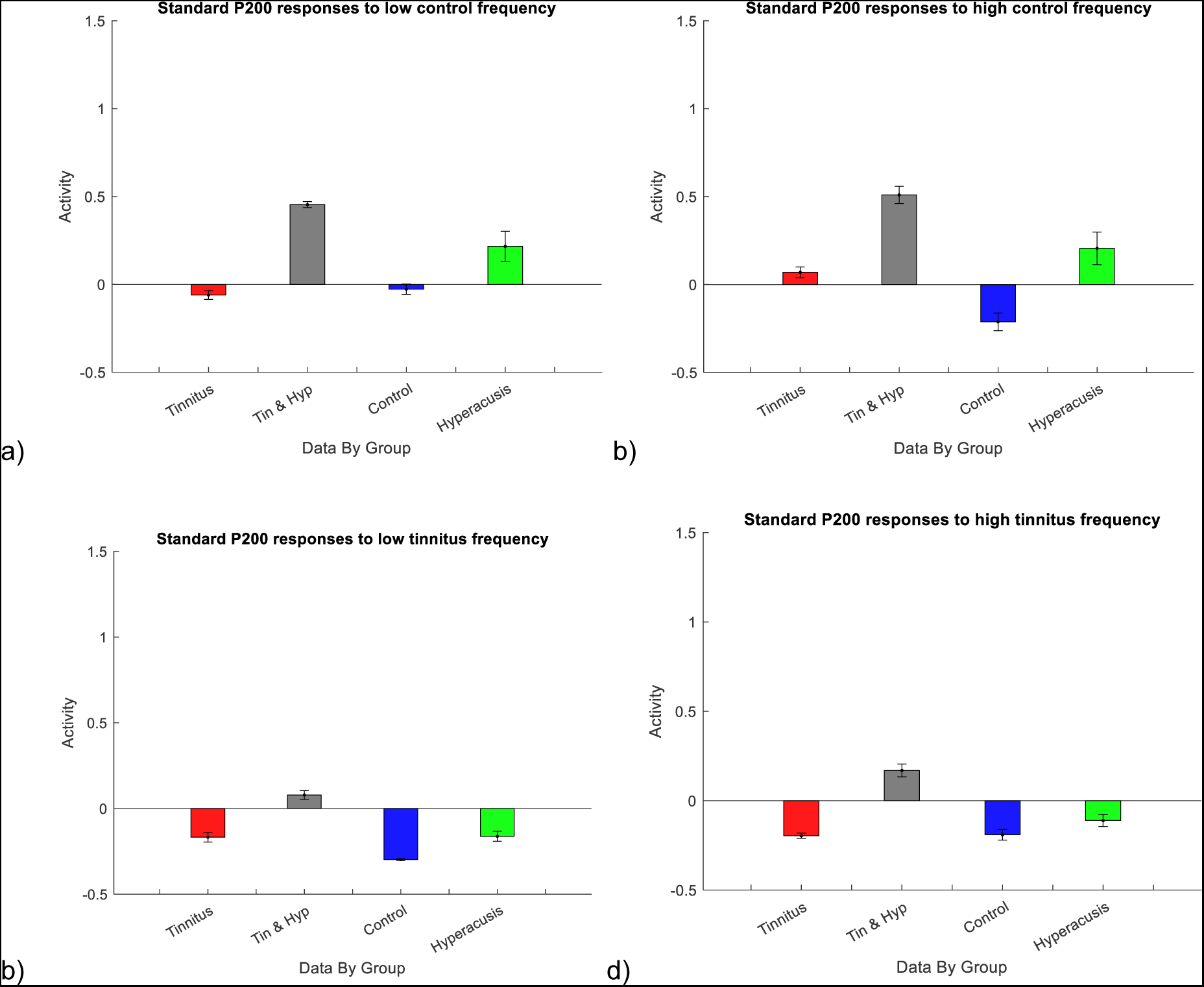
Standard responses in the P200 timeframe. Top two charts represent averaged responses to stimuli at 1 kHz frequency and bottom charts represent averaged responses to stimuli at tinnitus frequency. The order of the groups in all charts is: T+H- (red), T+H+ (grey). C (blue), T-H+ (green).

At the low intensity 1 kHz frequency, T+H+ group had significantly stronger p200 amplitude compared to all other groups (all p<0.001); T-H+ group had significantly stronger responses than T+H- and C groups (both p<0.001); no differences were seen between T+H- and C groups (p=0.948). At high intensity 1 kHz frequency, T+H+ group once again had significantly stronger responses than all other groups (p<0.001); C group had significantly lower responses than T+H- (p=0.002) and T-H+ (p<0.001) groups.

At the low intensity tinnitus frequency, T+H+ group had significantly higher responses than all other groups (all p<0.001) while C group had significantly lower responses than all other groups (all p<0.001). At the high intensity tinnitus frequency, again, T+H+ group had strongest responses (all p<0.001).

Overall, P200 responses to both high and low intensity standards were evident only in T-H+ and T+H+ groups at 1 kHz, and in the T+H+ group at the tinnitus frequency.

### Mismatch Negativity (MMN)

A difference waveform was calculated by subtracting standard waveforms from deviant waveforms, which was used for MMN analysis. Fig 8 shows MMNs in the 4 conditions. A three-way ANOVA (group, frequency, intensity) showed main effects of all factors, as well as significant interactions of all possible variations (all p<0.001). Post-hoc tests were split by frequency (fig 10).

**Fig 8.**
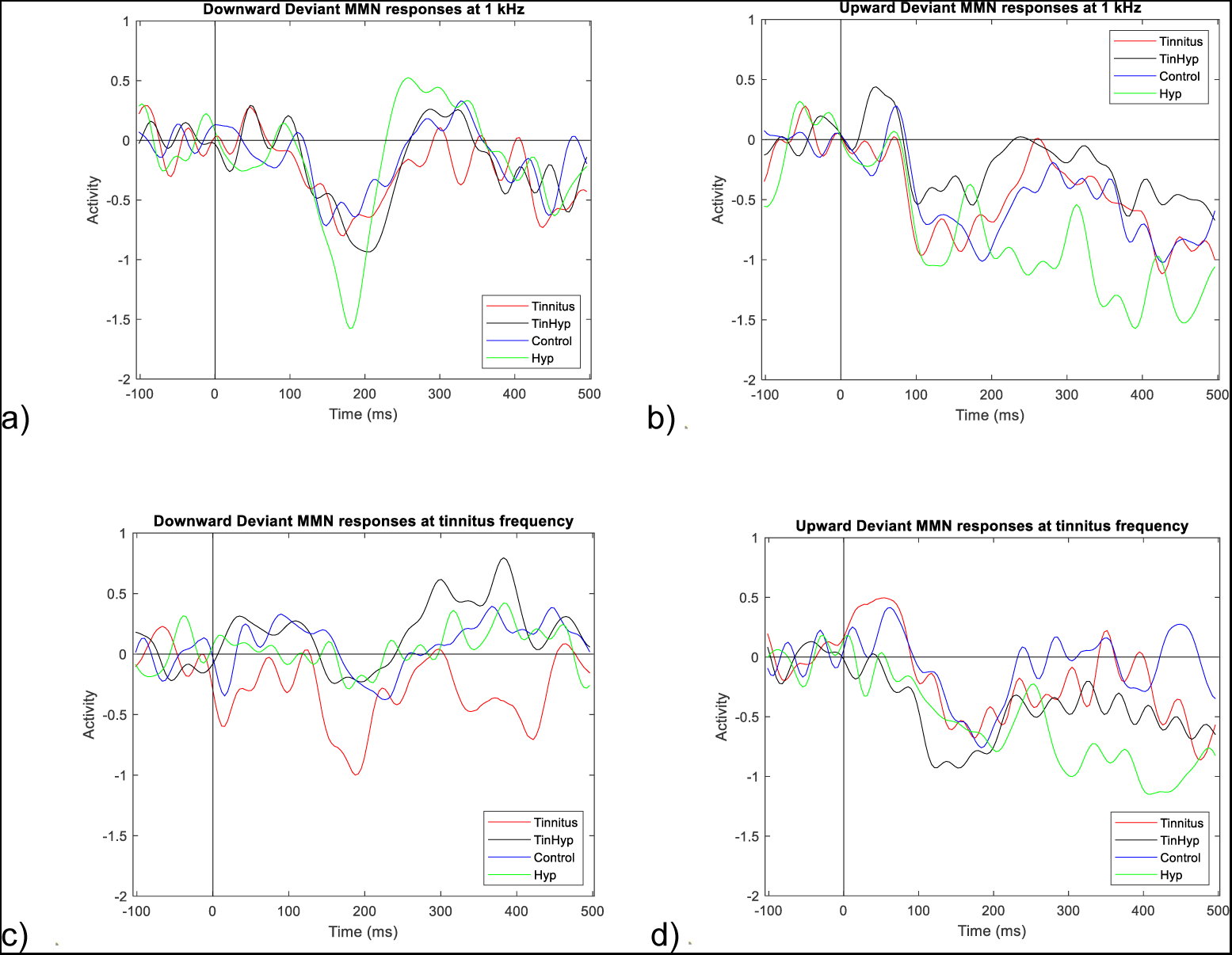
Average MMN waveforms for each subject group, in each condition. Graphs a-b show responses at 1 kHz frequency, and c-d show responses to the tinnitus frequency. On the left, are the downward deviants and on the right are the upward deviants.

**Fig. 10.**
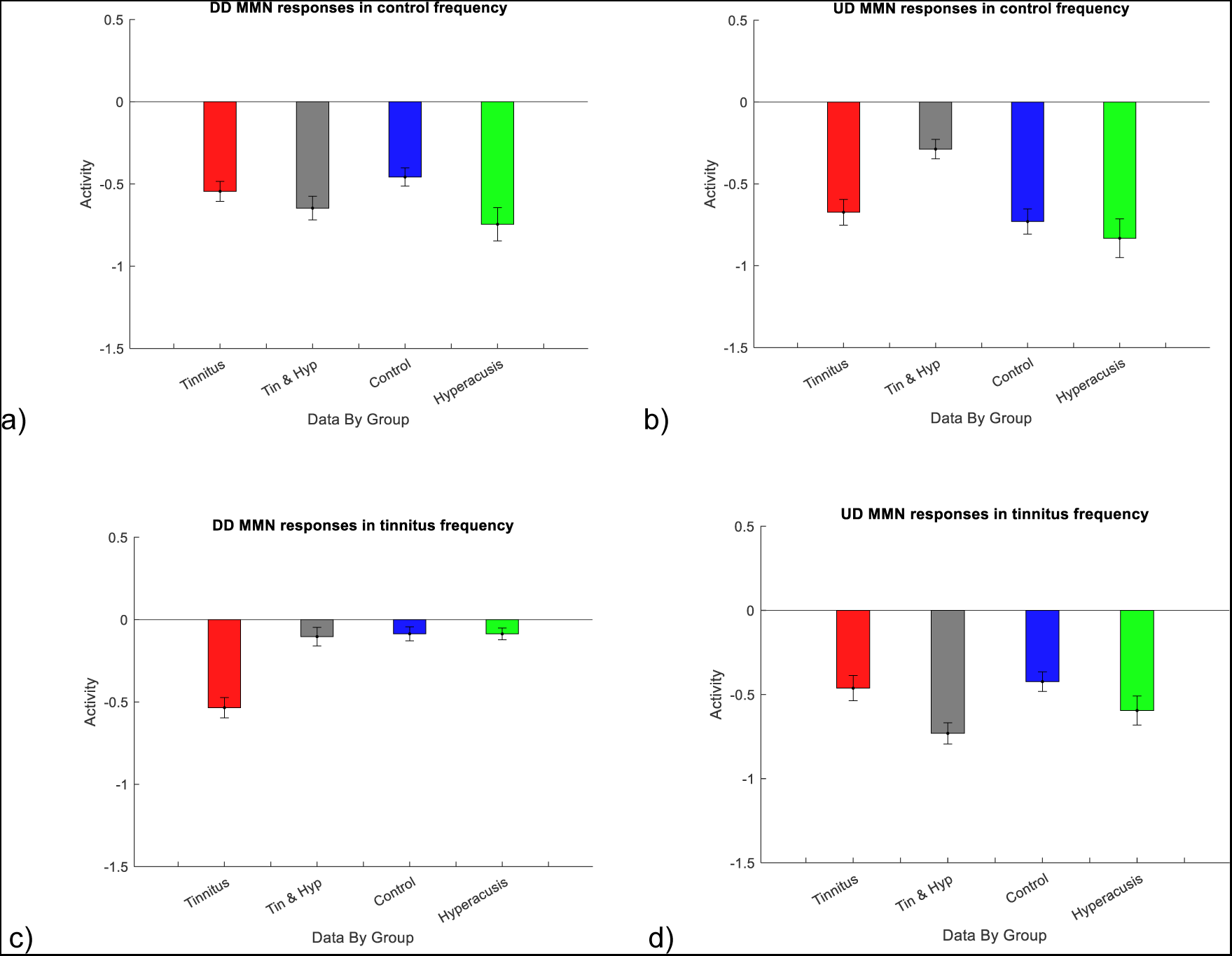
MMN responses. Top two charts represent averaged responses to stimuli at 1 kHz frequency and bottom charts represent averaged responses to stimuli at tinnitus frequency. The order of the groups in all charts is: T+H- (red), T+H+ (grey). C (blue), T-H+ (green).

At the 1 kHz frequency, T-H+ group had significantly stronger negative amplitude than the C group in response to DD (p=0.003), though there was also a trend towards a difference between T+H+ and C groups (p=0.064). In response to 1 kHz frequency UD, T+H+ group had significantly less negative amplitude compared to all other groups (all p<0.001).

At tinnitus frequency, responses to DD were significantly more negative in the T+H- group compared to all other groups (all p<0.001). In response to UD, T+H+ group had significantly more negative amplitudes compared to T+H- and C groups (p<0.001) and T-H+ group (p=0.024). T-H+ group also had significantly more negative amplitude compared to C group (p=0.002). There were no significant differences between T+H- group and C group (p=0.844).

### P50 timeframe difference waveform

The trend seen in the pure deviant p50 response (fig 2g), where T+H- group had significantly lower amplitude than the C group, may be present in the difference waveform as well, based on observation of fig 9. There, a two-way ANOVA (group, intensity) was carried out to see whether T+H- group would be significantly different from the other groups at tinnitus frequency specifically (Fig 11). The two-way ANOVA showed that both group and intensity had significant main effects (both p<0.001), and there was a significant interaction between group and intensity (p<0.001). Post-hoc Tukey tests showed that at DD difference waveform, T+H- had significantly lower amplitude than all other groups (all p<0.001); control group had no significant differences with either of the other groups (T+H+ p= 0.068, T-H+ p=0.098); however, there was also a significant difference between T+H+ and T-H+ groups (p<0.001).

**Fig 11.**
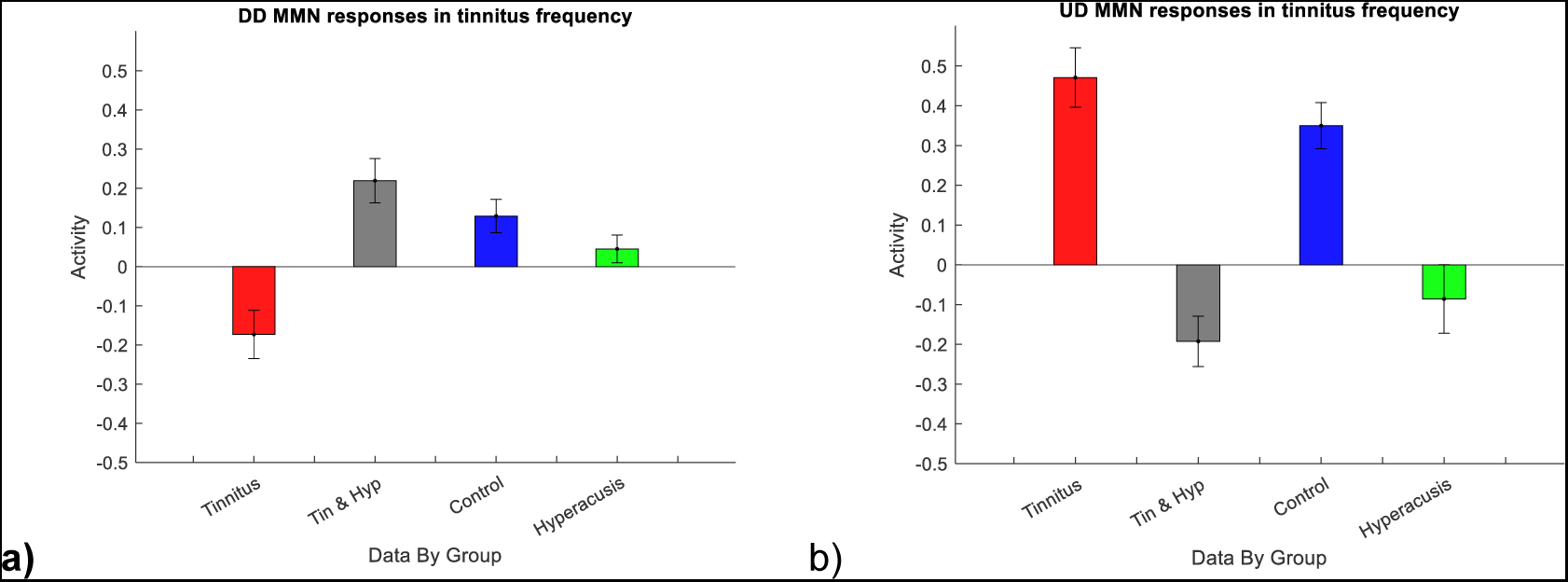
P50-timed deviant responses expressed as difference waveforms. The charts represent averaged responses to stimuli at tinnitus frequency. The order of the groups in all charts is: T+H- (red), T+H+ (grey). C (blue), T-H+ (green).

At UD difference waveform, T+H- group had significantly higher amplitude than all other groups (p=0.033 compared to C, p<0.001 compared to either of the H+ groups); T+H+ group also had significantly lower amplitude than the C group (p<0.001) but no significant difference compared to T-H+ group (p=0.071); C group also had significantly higher amplitude compared to T-H+ group (p<0.001).

## Results summary

Below are summary tables of each response type, comparing the 4 groups at each stage of the response (Table 2-4). The C group was likely representing the typical healthy response pattern. Based on these graphs, it appears that the MMN responses in T+H- group likely occurred due to the responses to the deviant tones. T+H+ group had the largest P200 responses in the standards and the deviants but also somewhat smaller P50s and larger N100 responses to the louder deviant tones, so it is possible that a mix of both tone types may be responsible for the altered MMN responses in this group. T-H+ group had some enhanced standard and deviant responses, so it is also possible that both tone types contributed to the MMN response.

**Table 2.**
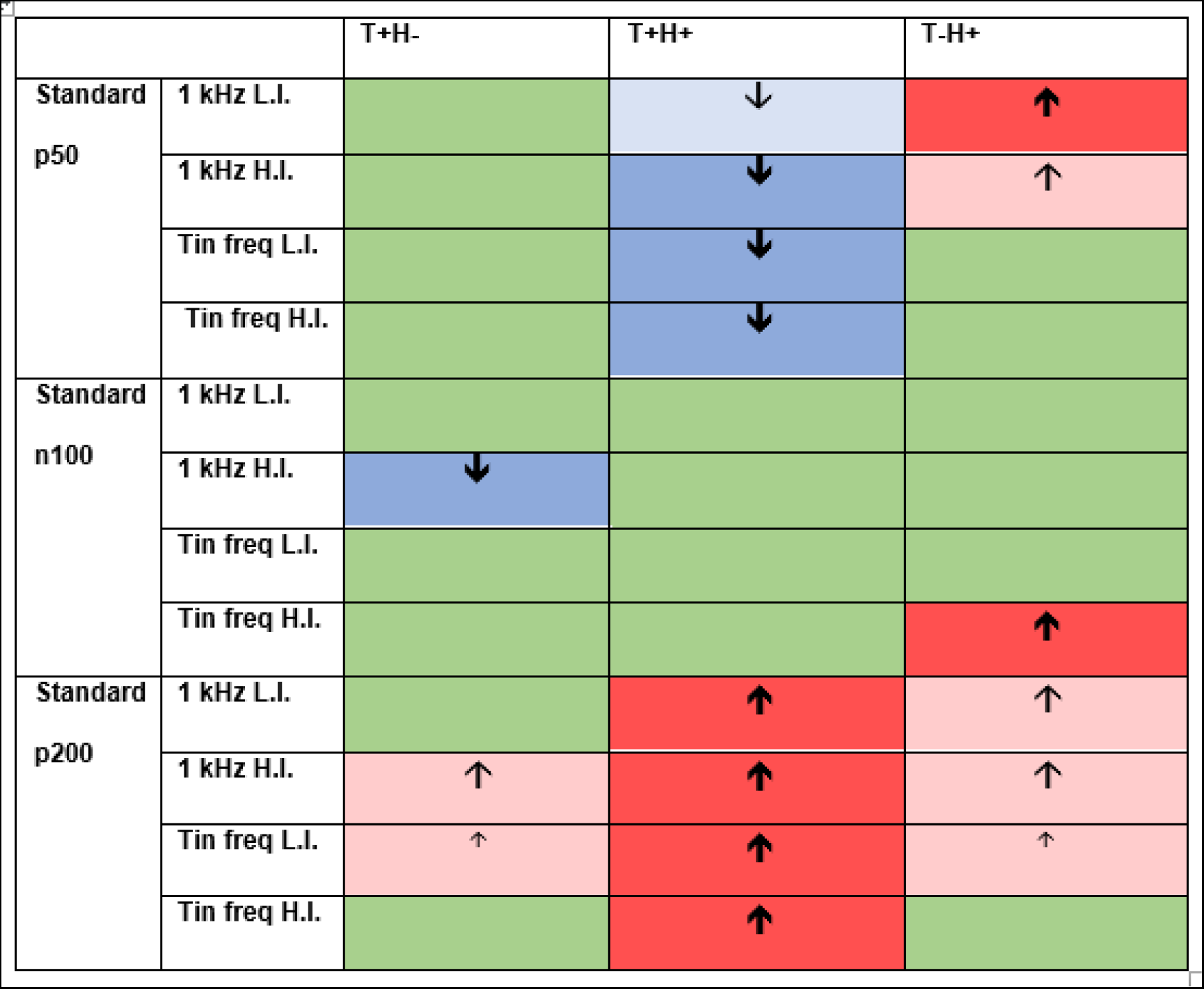
Summary of standard responses in groups with tinnitus and/or hyperacusis, compared to control group. Green indicates similar response amplitudes to C group; light blue indicates somewhat decreased amplitudes and darker blue with the thicker arrow represents most significant decrease compared to C group based on p-values. Light red represents increase compared to C group whereas bright red represent most significant increase. Tin freq = tinnitus frequency; L.I. = low intensity; H.I. = high intensity

|  |  | T+H- | T+H+ | T-H+ |
| --- | --- | --- | --- | --- |
| Standard<br>p50 | 1 kHz L.I. |  | ↓ | ↑ |
|  | 1 kHz H.I. |  | ↓ | ↑ |
|  | Tin freq L.I. |  | ↓ |  |
|  | Tin freq H.I. |  | ↓ |  |
| Standard<br>n100 | 1 kHz L.I. |  |  |  |
|  | 1 kHz H.I. | ↓ |  |  |
|  | Tin freq L.I. |  |  |  |
|  | Tin freq H.I. |  |  | ↑ |
| Standard<br>p200 | 1 kHz L.I. |  | ↑ | ↑ |
|  | 1 kHz H.I. | ↑ | ↑ | ↑ |
|  | Tin freq L.I. | ↑ | ↑ | ↑ |
|  | Tin freq H.I. |  | ↑ |  |

**Table 3.** Summary of deviant responses in groups with tinnitus and/or hyperacusis, compared to control group. Green indicates similar response amplitudes to C group; light blue indicates somewhat decreased amplitudes and darker blue with the thicker arrow represents most significant decrease compared to C group. Light red represents increase compared to C group whereas bright red represent most significant increase. Tin freq = tinnitus frequency; L.I. = low intensity; H.I. = high intensity

|  |  | T+H- | T+H+ | T-H+ |
| --- | --- | --- | --- | --- |
| Deviant<br>p50 | 1 kHz L.I. |  |  | ↑ |
|  | 1 kHz H.I. |  |  |  |
|  | Tin freq L.I. | ↓ | ↓ |  |
|  | Tin freq H.I. |  | ↓ | ↓ |
| Deviant<br>n100 | 1 kHz L.I. |  |  |  |
|  | 1 kHz H.I. |  |  |  |
|  | Tin freq L.I. | ↑ |  | ↑ |
|  | Tin freq H.I. |  | ↑ | ↑ |

**Table 4.**
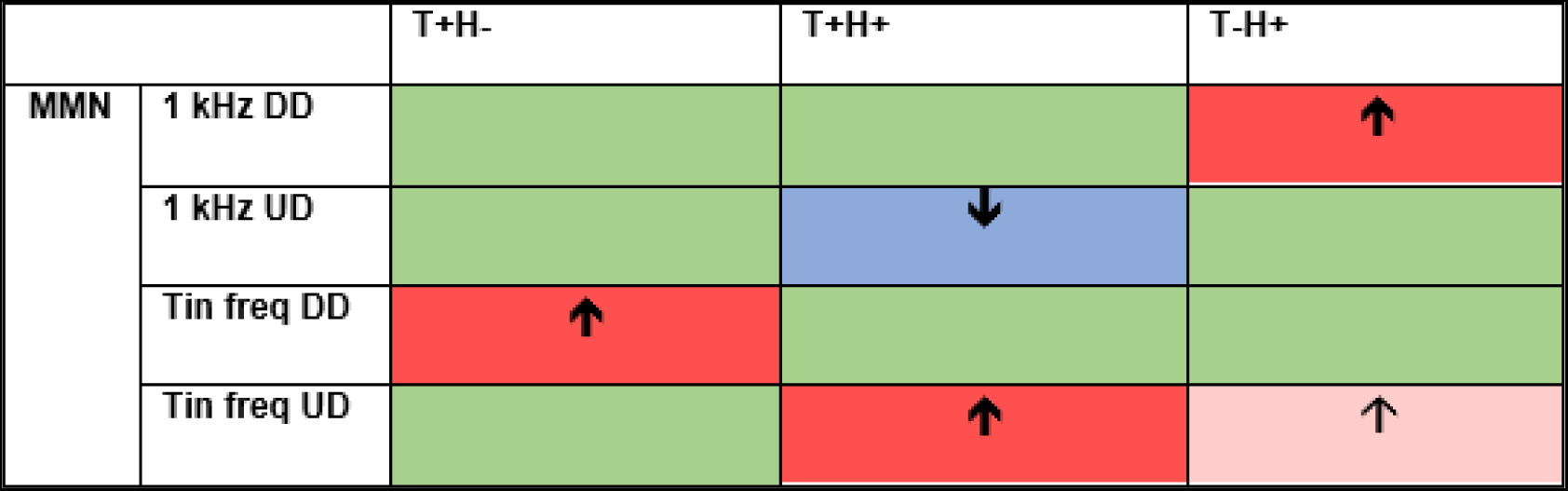
Summary of MMN responses in groups with tinnitus and/or hyperacusis, compared to control group. Green indicates similar response amplitudes to C group; light blue indicates somewhat decreased amplitudes and darker blue with the thicker arrow represents most significant decrease compared to C group. Light red represents increase compared to C group whereas bright red represents the most significant increases. Tin freq = tinnitus frequency.

|  |  | T+H- | T+H+ | T-H+ |
| --- | --- | --- | --- | --- |
| MMN | 1 kHz DD |  |  | ↑ |
|  | 1 kHz UD |  | ↓ |  |
|  | Tin freq DD | ↑ |  |  |
|  | Tin freq UD |  | ↑ | ↑ |

## Discussion

The current study shows distinct profiles for tinnitus and hyperacusis, as well as additional more nuanced interactions, which not only moves our understanding of each condition, but also speaks directly to possible mechanistic subtypes of tinnitus (and of hyperacusis) which might be disentangled through the cheap and available technique that is single-channel EEG. The current findings may explain some discrepant findings in past literature.

### Summary of results

#### Correlates of tinnitus with/without hyperacusis

There were some distinct differences seen between the response patterns of T+H+ and T+H- groups. In the presence of hyperacusis in the T+ groups, p50 amplitudes were smaller in response to standard tone of 1 kHz at high intensity and in response to standard tone of tinnitus frequency at low intensity. Additionally, though both T+ groups had weaker P50 responses to raw deviant tones at tinnitus frequency, the two groups had slightly different patterns. T+H+ participants had stronger responses to low intensity stimuli but weaker responses to high intensity stimuli compared to T+H- group (T+H+ group seemed to involve both differences seen in T+H- and T-H+, therefore potentially showing a non-direction-specific reduction in pre-attentive change detection at tinnitus frequency [53]). Usually, P50 is suppressed in response to a repeating stimulus. Therefore, the smaller P50 amplitude may indicate a greater degree of sensory gating in tinnitus groups, particularly around the tinnitus frequency. Notably, in T+H+ even the P50 to standards are decreased compared to C group, in all conditions.

The difference in P50 amplitude remained significant in the difference waveform. There was an opposing pattern between the two T+ groups, as well as a difference with the C group. This could not be explained by higher tinnitus severity as previously suggested, as the decreased p50 (and following more negative N100) occurred in both T+ groups just in different conditions, though heightened cognitive awareness may be playing a part [46]. In the T+H- group, P50 for UD is similar to C group responses, however, P50 is significantly diminished for DD. It is possible that this occurs due to suppression of ascending input when the sensory input first decreases, but it could also indicate a lower reliance on sensory input and a greater reliance on higher predictions compared to healthy participants. In a previous paper, it was suggested that participants with tinnitus could be less able to attend to relevant stimuli due to underactive feedback inhibition in the auditory microcircuits specifically in the tinnitus frequency regions [54], which may lead tinnitus subjects to over-rely on higher predictions in response to diminished sensory input. However, this also means that they cannot ignore UDs due to hyper-excitability and therefore respond with higher alertness to UDs. On the other hand P50 in T+H+ are greatly suppressed in response to UDs, but not to DD, in a similar fashion to T-H+, therefore this may be an effect of hyperacusis.

N100 had almost an opposite pattern to raw p50 responses. The presence of hyperacusis was related to stronger response to high intensity standard tones at 1 kHz, weaker response to low intensity deviant tones at tinnitus frequency, as well as stronger response to high intensity deviant tones at tinnitus frequency. Potentially, this process allowed the two groups to even out sensory gating in the pre-attentive phase prior to MMN/P200 timeframes. MMN responses somewhat followed the N100 deviant response pattern at the tinnitus frequency: DDs elicited larger responses in T+H- group, whereas T+H+ group had stronger response to UDs. Additionally, T+H+ group had weaker responses to upward deviants at 1 kHz tone. In tandem to the potential theory about P50s, it is possible that for T+H-, stronger predictions would have greater prediction error responses when not completely fulfilled (i.e. in DDs, which can be thought of as incomplete omission responses). However, for the H+ groups the stronger MMN response to tinnitus frequency UD likely occurred due to heightened perception of changes in intensity.

### Comparison to previous studies

A previous study mentioned in the introduction found stronger subcortical sound-evoked responses in T+H+ participants compared to T+H- [11]. This finding was not frequency-specific, but it is corroborated by our data: there was increased sensory gating at P50 (reduced amplitude) but a heightened P200. Additionally, the previous research showed that at 1 kHz, an UD increased N100 and P200 amplitudes in bilateral T+ group compared to C group. In our study, while P200 was increased in both groups in response to a 1 kHz UD, N100 remained similar to C group. This could be due to a 20 dB SPL rise in the previous study, compared to average of 6 dB in the current study, but the trend of larger responses can still be seen, especially in T+H+ group. However, the current study found that in response to stimuli at tinnitus-like frequency, the T+H+ group showed significantly smaller activation than T+H- group [11]. Importantly, in the aforementioned previous study, tones used were loudness-matched tones at frequencies ranging between 250 Hz to 8 kHz, with the most tinnitus-like tone used for this comparison, therefore the previous findings may not have been truly be representative of a response to a tinnitus-like frequency. This is an interesting divergence compared to the current findings, which could be due to design and more precise tinnitus frequency matching.

A multi-feature paradigm study mentioned in the introduction [49] concluded that people with high noise sensitivity showed significantly diminished P50s to all deviants and significantly less negative MMNs to noise deviants (when corrected for multiple testing) than the low noise sensitivity group. The pattern of the T-H+ group in the current study did not follow the previous finding, but T+H+ pattern of P50 responses did. However, the reason for this could be the more complex nature of the presented sounds (chords) as well as potentially different causes of hyperacusis in different groups.

Unfortunately, it is not possible to make frequency comparisons as these researchers did not explicitly state the frequencies at which tones were played (though from figures, base tone seemed to be around 2 kHz). While not directly related the deviant used in the current study, this paper was one of the few studies investigating MMN and noise sensitivity specifically. Further, a number of multi-feature deviant paradigm papers reported that generally, MMN responses in tinnitus participants for all deviant types used tended to be smaller (e.g. [53, 55, 56]), both at higher and lower frequencies.

Studies that focused on 0.5, 1 and 1.5 kHz frequencies saw that tinnitus presence was related to smaller amplitudes for a number of deviant types (though intensity deviants were inconsistent in showing any difference in amplitudes in tinnitus compared to control groups, and some studies not controlling for hyperacusis, while studies that did not show a difference being similar in their finding to our T+H- and C group MMN responses to 1 kHz frequency deviants in the current study) [50, 53, 57]. A number of studies have also looked at higher frequencies, e.g. 5 kHz and 8 kHz ([56, 57]). While at 5 kHz, once again, tinnitus presence was related to weaker MMN amplitudes [57], at 8 kHz a study found that when a participant was habituated to their tinnitus, responses were similar to controls, but when the participants were not habituated to their tinnitus, MMN amplitudes were weaker (for higher frequency deviant) [56]. No such findings were seen in the current study, rather an amplitude increase was seen at the tinnitus frequency. It is possible that the differences are due to a different deviant type (intensity vs. frequency) or because not all participants with tinnitus would find 8 kHz to be near the tinnitus frequency. Here, it was found that at 1 kHz, T+H- group had similar responses to C group, while T+H+ group had a weaker MMN response to UD at 1 kHz. T+H+ group also had higher THI scores so possibly overall the group would be less habituated to their tinnitus than T+H- group. On the other hand, it is possible that diminished auditory responses in such paradigms may reflect an auditory dysfunction more generally, or may reflect presence of noise sensitivity rather than tinnitus, or any mixture of the two conditions. The current study did not use a multi-feature paradigm which may be the reason why the current study did not always see similar patterns. Rather than seeing the general effect of auditory dysfunction, we were able to disentangle alteration related to different conditions as well as their combined effects. The current study was rigorously controlled for hearing loss and hyperacusis presence, which may contribute to the disparities with previous findings.

#### Correlates of hyperacusis with/without tinnitus

There were significant distinctions between T+H+ and T-H+ groups, in relation to each other as well as in comparison to C group. Presence of tinnitus in people with hyperacusis seems to have the opposite effect on ERP amplitudes compared to people only with hyperacusis at P50 and standard N100. T-H+ group had highest amplitudes to all P50 standards (though only significantly higher at 1 kHz), while T+H+ responses were weaker than both H- group responses at all frequencies, which may indicate opposite sensory gating alterations between the groups. T-H+ also had the highest P50 amplitude to low intensity deviant at 1 kHz, and strongest N100 in response to both deviant conditions at tinnitus frequencies, potentially indicating lack of habituation. N100 has previously been shown to be suppressed during repetitive stimulation, through a process called repetition suppression (RS) [58]. The N100 deviant responses at tinnitus frequency in T+H+ were lower than both H- group responses, though not significantly. T-H+ also had stronger N100 response to high intensity at tinnitus frequency standards, also indicating lack of habituation to a high intensity, potentially uncomfortable tone.

However, this opposing trend between the groups changed in the later ERPs. P200 response at both frequencies was higher in T+H+ than T-H+, though T-H+ was still stronger than both H- groups at 1 kHz. The result was difficult to interpret at high intensity tinnitus frequency standard, as T-H+ P200 response followed a significantly lower N100 and still visually had a peak around 200 ms but the inferential statistics saw this as non-significant. At 1 kHz, MMN responses were strongest to DD in T-H+ and weakest to UD in T+H+. At the tinnitus frequency, strongest response to UD was in T+H+ group, followed by T-H+. So, presence of hyperacusis reduced P50 in T+ participants but increased P50 in T-participants, however, later in the timeframe, both groups show an enhanced P200 response to both standard and deviant tones at tinnitus frequency, particularly in the T+ group (despite T+H- group showing amplitudes closer to controls).

Comparison to the original IMA study is slightly limited, as the the earlier paradigm included an edge frequency of tinnitus (sound just below the tinnitus frequency of a participant), but not a 1 kHz control frequency [24]. However, the later follow-up study included 1 kHz condition. The tinnitus group in the previous studies had non-significantly smaller amplitudes at tinnitus frequency. This is similar to the findings in the current study, where the two T+ groups were mostly similar to C group, but the T+H+ group had lower p50 amplitudes especially at tinnitus frequency while T+H- group had less strong n100 at 1 kHz in response to high intensity standards. If the two groups were mixed, it likely would reduce their differences with C group and would corroborate the previous findings. Overall, T+H- participants did not follow the original study pattern. However, the T+H+ group of the current study had a very similar pattern to original study and the later follow-up study in the edge tinnitus frequency. The current results strongly indicate that previous differences seen previous were driven by hyperacusis rather than tinnitus [25]. Additionally, C group patterns seem similar in the current study and the earlier follow-up study, but different in the original study, which may be due to overall frequency differences in the paradigm.

#### Comparison to previous findings

In a paper comparing responses of controls and subjects with low vs high tinnitus-related distress to three tone bursts (1 kHz, 1.3 kHz and 1.6 kHz at 90 dB HL), participants either had to press a button when they heard a particular tone, or ignore the tones [59]. Findings showed that unattended stimuli elicited weaker N100 amplitudes than attended stimuli, which was only significant in the low tinnitus distress group, but not high distress. Additionally, high distress tinnitus group had larger overall N100 amplitudes than either low distress or (especially) controls, indicating potentially reduced ability to habituate to the repetitive stimuli. Researchers concluded that high tinnitus distress is related to more attention paid to the tinnitus; the mean threshold of uncomfortable loudness was also similar between the groups in both ears, so presence of hyperacusis was unlikely to explain the results. Interestingly, in the current study, a consistent trend for weaker N100 (significant for high intensity, non-significant trend for low intensity) was seen in T+H- group at 1 kHz compared to C and T+H+ group. This was also the case at tinnitus frequency low intensity, but not high intensity standards. So, the T+H- group either exhibited the most habituation, out of the four groups, shown by ability to suppress the N100 response, in all tones except for the sound of their tinnitus becomes louder. This could also be potentially explained by an overall tendency of T+H- participants for higher reliance on higher predictions in the auditory system. While the previous study did not test T-H+ participants specifically, in the current study it seems that the T-H+ were the least able to suppress the N100 to repeating tones in all conditions.

#### Repetition positivity & habituation

Repetition Positivity (RP) is an ERP that has been established as a function of memory trace formation and potentially suppression of prediction errors (opposite from MMN), which presents as a positive wave around 200 ms that is enhanced by stimulus repetition [58, 60]. A systematic review of the P200 ERP has suggested that P200 amplitude is linearly related to increase in intensity of an auditory stimulus, and may even reach a saturation level at high intensities [33]. Additionally, P200 presence may relate to working memory and interference control but prolonged P200 latency may reflect early conscious attention towards a certain stimulus [61]. This may be interesting as the T+H+ group had longer P200 latencies than T-H+ group in response to 1 kHz stimuli, while T-H+ group tended to have an earlier peak within the P200 timeframe. Therefore, it is possible that H+ participants generally either perceive the stimulus as louder than H- participants and therefore have higher amplitudes of P200, or they use more interference control. T+H+ in particular may also pay more conscious attention to the stimuli as their P200 lasts longer than in T-H+ group. Perceiving the higher intensity as louder than H- groups may also in part explain the increased MMN response to UD in both H+ groups at the tinnitus frequency, and the increased conscious attention could explain why MMN response to UD was particularly high in T+H+. However, it could not explain the MMN response pattern at 1 kHz (increased amplitude in T-H+ to 1 kHz DD and decreased amplitude in T+H+ to 1 kHz UD). However, there is also a possibility that these differences were simply due to lower alertness in the H- groups, as lower P200 responses have previously been seen in healthy participants in ‘mind wandering’ condition during an oddball task compared to ‘focused on breathing’ condition [62].

### Future directions

Taken together, the findings indicate that tinnitus researchers need to account for hyperacusis when utilising ERP-based paradigms, and research into correlates of hyperacusis must not focus solely on tinnitus-associated hyperacusis, especially because tinnitus and hyperacusis combined produces different patterns compared to people with only one of the conditions. Additionally, there may be a need to understand whether the frequencies used in the paradigm affect the patterns that are seen among groups, including controls. For example, the pattern of MMN responses was different in the original study compared to the replication study. An MEG study investigated repetition positivity alongside repetition suppression, in which they established that the two processes are separate but complementary. N100 was suppressed during repetitive stimulation, while an increasing later field was seen following repetition of a stimulus [58]. An interesting suggestion was made regarding two separate RP generators, one located more frontally (Fz) that is dependent on longer-term paradigm contexts (one tone vs roving paradigm) and another located towards the mastoids and dependent on short-term changes [63], which potentially affects overall presentation of RPs [64]. This further calls for investigation of ERP patterns in different global contexts, as well as potential further study of standards in tinnitus and/or hyperacusis, as these conditions seem to affect RS and RE processes in different ways. Determining the expected patterns in varying overall paradigm contexts would allow researchers to better understand how activity is altered in the presence of tinnitus and/or hyperacusis.

### Limitations

A potential limitation of this study is that there are a number of types of hyperacusis (e.g. loudness vs pain subtypes), as well as prevalence of autism-related symptoms or comorbid pain disorders in people with hyperacusis, which may all have differing effects on brain activity [5, 65, 66]. Additionally, it may be useful in the future to conduct a source analyses-based study in order to investigate how the presence of hyperacusis and tinnitus work together, compared to presence of one of the conditions only.

### Conclusion

Hyperacusis has quite a broad influence on evoked responses, including early and late, and standard and deviant, across frequencies. For example, it is related to larger P50s and P200s, and particularly increased MMN responses to UDs. Overall, this indicates a hypersensitivity to unexpected intensity increases. With presence of tinnitus and hyperacusis, it seems that the later responses align with, and are potentially enhanced by, the patterns seen with hyperacusis alone. However, the earlier (P50) responses are suppressed, possibly exaggerating the effect of tinnitus alone on the evoked responses. Additionally, hyperacusis seen in tinnitus seems to have effects that are more limited to the tinnitus (hearing loss) frequency, unlike hyperacusis without tinnitus. This could be due to slightly different mechanisms between the two types. Tinnitus itself has a more specific pattern, almost limited to the tinnitus frequency only, with most of these evoked responses being normal (though, a few differences such as smaller P50 responses to standards, and a distinct pattern of P50 changes in deviants: smaller to downward and larger to upward), but a strikingly different (larger) MMN to downward intensity deviants at the tinnitus frequency only. This is potentially indicative of stronger formations of auditory predictions or memory traces around the tinnitus frequency (the area affected by hearing loss).

